# Hybrid Cellular Automata Modelling Reveals the Effects of Glucose Gradients on Tumour Spheroid Growth

**DOI:** 10.1101/2023.10.19.563082

**Authors:** Luca Messina, Rosalia Ferraro, Maria J. Peláez, Zhihui Wang, Vittorio Cristini, Prashant Dogra, Sergio Caserta

**Author notes:** Correspondence: Sergio Caserta, or Prashant Dogra.

## Abstract

**Purpose:** In recent years, mathematical models have become instrumental in cancer research, offering insights into tumor growth dynamics, and guiding the development of pharmacological strategies. These models, encompassing diverse biological and physical processes, are increasingly used in clinical settings, showing remarkable predictive precision for individual patient outcomes and therapeutic responses.

**Methods:** Motivated by these advancements, our study introduces an innovative *in silico* model for simulating tumor growth and invasiveness. The Automated Hybrid Cell emulates critical tumor cell characteristics, including rapid proliferation, heightened motility, reduced cell adhesion, and increased responsiveness to chemotactic signals. This model explores the potential evolution of 3D tumor spheroids by manipulating biological parameters and microenvironment factors, focusing on nutrient availability.

**Results:** Our comprehensive Global and Local Sensitivity Analyses reveal that tumor growth primarily depends on cell duplication speed and cell-to-cell adhesion, rather than external chemical gradients. Conversely, tumor invasiveness is predominantly driven by chemotaxis. These insights illuminate tumor development mechanisms, providing vital guidance for effective strategies against tumor progression. Our proposed model is a valuable tool for advancing cancer biology research and exploring potential therapeutic interventions.

**Simple Summary:** In recent years, mathematical models have revolutionized cancer research, illuminating the complex dynamics of tumor growth and aiding drug development. These models, reflecting biological and physical processes, are increasingly used in clinical practice, offering precise patient-specific predictions. Our work introduces an innovative in silico model to simulate tumor growth and invasiveness. The Automated Hybrid Cell, replicating key tumor cell features, enables exploration of 3D tumor spheroid evolution. Sensitivity analyses reveal that tumor growth is primarily influenced by cell replication speed and adhesion, while invasiveness relies on chemotaxis. These insights shed light on tumor development mechanisms, guiding effective strategies against tumor progression. Our model serves as a valuable tool for advancing cancer biology research and potential therapeutic interventions.

## 1. Introduction

According to the World Health Organization, cancer remains one of the leading causes of death in advanced countries. In 2020 alone, there were 19.3 million new cases and 10 million deaths reported worldwide [1,2]. However, there has been a consistent decrease in cancer-related mortality, primarily due to successful prevention efforts and advancements in treatment options [3]. It is noteworthy that approximately 66.7% of cancer deaths are related to a process known as metastasis [4], which refers to the formation of secondary tumours in parts of body different from the site of cancer origin. The process involves cancer cells crossing endothelial walls, circulating through in the bloodstream, and eventually extravasate from capillaries crossing the basement membrane to colonize the new site [5].

A key feature in the above steps is the ability of cancer cells to migrate directionally in response to external stimuli, known as chemotaxis, which plays a fundamental role in enabling tumour invasiveness [6]. Chemotaxis is defined as the directional movement of an organism in response to the concentration gradient of a given chemical species. In the context of cell chemotaxis, it refers to the sensitivity of cells to specific gradients, quantified as the ratio between the concentration gradient and the local concentration value (S = ∇C/C) [7,8]. Chemokines (e.g., interleukin) and growth factors (e.g., FBS) are examples of chemical species that drive chemotaxis in cancer cells [7]. Metabolites such as glucose or oxygen, which are also present in many growth factors, as well as catabolite gradients, can also induce chemotaxis.

In the case of cancer tissues, cell over-proliferation induces a lack of nutrients and an accumulation of catabolites. Consequently, tumour masses lose compactness and, in order to keep a high surface to volume ratio, tend to invade surrounding tissues. This morphologic instability [9], known as diffusional instability [10,11], can be predicted by mathematical models. The investigation of such a complex phenomenon requires the use of adequate models able to include both biological and transport phenomena aspects. Different approaches have been followed in cancer research, ranging from simple 2D cell cultures, complex 3D scaffolds *in vitro*, murine *in vivo* models, up to clinical studies.

In cancer research, *in vitro* approaches, which involve the study of biological systems in artificial laboratory conditions, have been widely employed to investigate the underlying mechanisms of cell migration, proliferation, and other processes related to tumorigenesis. Most of the published data regarding known cell-based processes are derived from experiments performed in two-dimensional (2D) conditions. 2D cell cultures, growing on solid substrates such as plastic, do not fully reproduce the complex three-dimensional (3D) architecture and complexity of living tissues. Important aspects such as cell-cell interactions, tissue phenotypes, the role of cell density and extracellular matrix (ECM) [12-15], proliferation regulators [16], and metabolic functions [17] are often missing or not adequately represented in 2D cultures.

In order to mimic the native *in vivo* scenario [18-20] where cancer growth happens, 3D models have been used in cancer research as a compromise between 2D cell cultures and whole-animal systems. However, 3D tumour microenvironments in murine models may not accurately represent the human scenario, making it challenging to control and interpret experimental outcomes.

Recently, tumour spheroid [8], a tightly bound aggregate of cells, is gaining popularity as 3D model due to of its ability to strikingly mirror the 3D cellular context. Being characterized by naturally physiological and chemical gradients, cellular spheroids consist of actively proliferating cells on the outside with quiescent cells in the inner rim, and a central nutrient deprived zone (necrotic core) [21], mimicking the natural scenario in a tumour *in situ*.

Each of the above-mentioned approaches has limitations, that can be taken into account by adequately coupling experimental investigations with *in silico* mathematical models. One of the main challenges is to capture the complex and dynamic interactions between cancer cells and their microenvironment. In vitro experimental conditions may introduce artifacts that can potentially compromise the validity of conclusions drawn from these studies if not properly accounted for. *In silico* approaches, based on mathematical modelling and computer simulations, have the potential to overcome these limitations by focusing on the representation of the interactions of cells with microenvironment, that are hard to mimic experimentally [22,23]. Furthermore, *in silico* models, once developed and adequately validated, can be used to conduct extensive virtual experimental campaigns at a significantly lower cost compared to *in vivo* and *in vitro* research. This approach is crucial for the fundamental understanding of mechanisms of cancer growth and can facilitate the simulation of specific treatments on individual patients, or the study of the effects of rare mutations or genetic variations, implementing precision and personalized medicine [24] protocols.

For this reason, *in silico* approaches [25] are attracting growing attention. The earliest attempts to mathematically model tumour growth and invasiveness dates back to 1967, and since then, the number and accuracy of these models have continued to grow [14]. These models can be classified into continuum models, discrete models, and hybrid models [26] and they are employed to simulate and study a wide variety of phenomena, such as the resistance of tumours to different drug treatments [27-29], the interaction between tumour cells and their microenvironment [13,15] and the immune system [25,30].

Hybrid agent-based models, such as Hybrid Cellular Automata (HCA), have proven to be effective in investigating complex tumour systems by combining deterministic reaction-diffusion partial differential equations with the representation of the cells as single and autonomous entities (Cellular Automata, CA [25]) governed by deterministic or stochastic laws. These models consider interactions between individual cells[26-29,31] and can provide deeper insights into the mechanisms underlying cancer growth and progression, supporting the design of new therapeutic strategies [30,32].

HCA models are particularly useful in many applications in cancer research, where a multi-scales model required to depict the dynamics of cancer development over time, including the evolution of cell phenotypes and genotypes [25]. These models account for the interaction of individual cells with parameters characterizing the surrounding environment, such as concentration fields or changes in pH [33].

In this study, a computational model based on HCA was developed to simulate the growth and invasiveness of spheroids under different gradients of nutrients. Specifically, chemotaxis was taken into account by estimating glucose concentration profiles surrounding a cancer cell aggregate (tumour spheroid). The model aims to investigate the role of cell migration, duplication, and other parameters such as cell-cell adhesion and sensitivity to chemotactic gradients in the phenomenon of cancer invasion. A Global Sensitivity Analysis (GSA) and a Local Sensitivity Analysis (LSA) were performed to evaluate the model’s sensitivity to variations in the above-mentioned cellular parameters.

The goal of this study is to develop a model that can capture the complexity of the cancer invasion mechanism, as envisioned by the diffusional instability model [9], while maintaining computational feasibility. This will provide a more comprehensive understanding of cancer biology and aid in the design of new therapeutic strategies.

## 2. Materials and Methods

In the following section the structure of the 2D-HCA model used for simulating *in vitro* growth of avascular tumour spheroids will be presented in detail. The model description is organized into several subsections. Firstly, the model domain (2.1.1) and numerical methods used to calculate continuous functions (in our case glucose concentration) (2.1.2) are presented. HCA model will be described presenting cell phenotypes (2.1.3) and rules governing cellular dynamics (2.1.4). Statistical and sensitivity analysis will be described in an independent subsection (2.2).

By structuring the model description in this manner, the paper provides a comprehensive overview of the 2D-HCA model and its application in simulating the in vitro growth of avascular tumour spheroids.

### 2.1. Model development

The 2D domain used in the model represents a layer of the ECM that contains a single tumour spheroid. This domain is discretized into a squared grid, referred to as a lattice. Each element of the lattice (lattice point, LP) can be occupied by a single cell or remain empty, representing the presence of ECM only. The entire grid evolves through a series of discrete time steps, following a set of pre-defined rules. The evolution of each cell is governed by the chemical stimuli in the LP occupied, and by the state of neighbouring LPs. In this model, the chemical stimuli are represented by the concentration profile of a chemoattractant, specifically glucose. The model can be viewed as a superposition of two identical square grids, where one represents the cellular layer (see **Cellular Layer** section) and the other is used to calculate the glucose concentration field (see **Glucose Layer** section). The model can be easily generalised to account for more chemical species, or other type of stimuli such as pressure fields, simply adding further layers.

In **Figure 1a**, a schematic representation (not in scale) of the cellular layer superimposed to the glucose layer of our model is presented. In the cellular layer, the LPs are coloured either in orange or blue, corresponding to positions occupied or not by cells. In each LP, glucose concentration is calculated and schematically represented in **Figure 1a** in a colour scale. As qualitatively reported in **Figure 1a**, glucose concentration at the grid margins is higher compared to the centre of the domain. The resulting symmetrical radial concentration gradient is due to an isotropic source of glucose from each of the four edges of the domain, and a consumption in the centre where cell spheroid is located.

**Figure 1.**
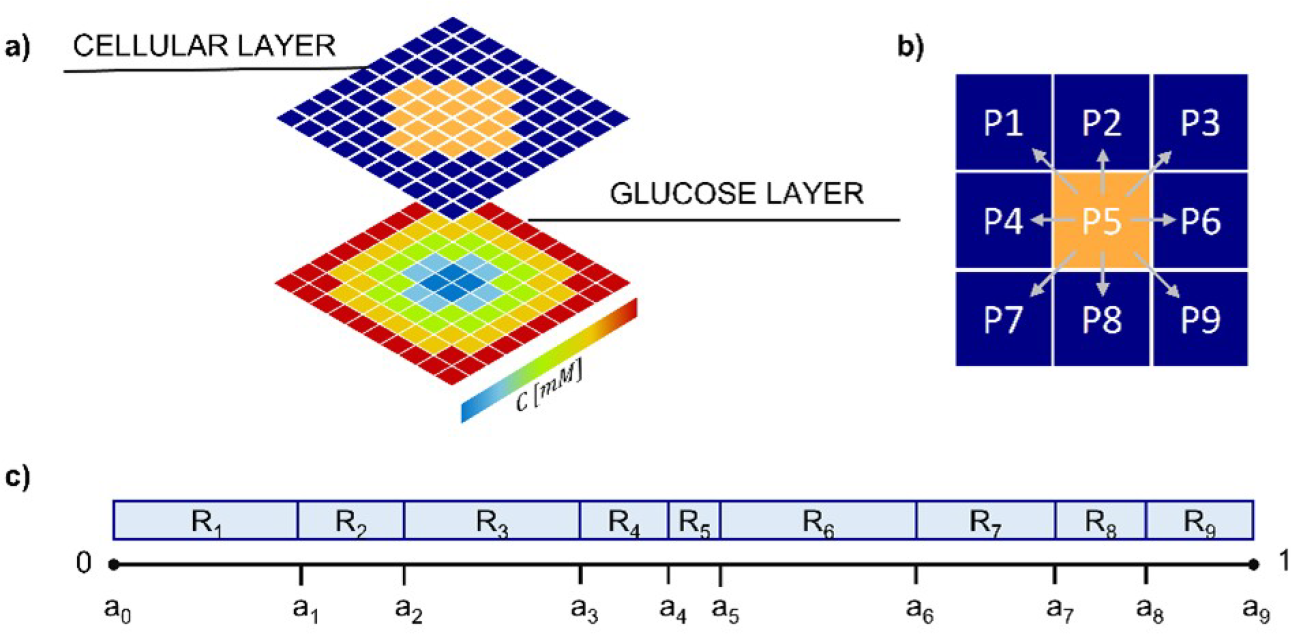
Model schematic showing cellular and glucose field layers. **a)** Typical initial configuration of the Cellular Layer (upper grid) with cell-free LPs (dark blue) surround cell-occupied LPs (orange). Not in scale. In the Glucose Layer (bottom grid) glucose concentration is calculated in pseudo-stationary condition in each Lattice Point (LP), according to Eq. 6. In the scheme a typical concentration profile in a cell spheroid is reported using a colour scale. **b)** 8 possible migration directions for a representative cell (located in P5); **c)** Schematic representation of the migration direction probabilities (R_i_ values).

#### 2.1.1. Domain building

In this study, the domain considered for the model is a squared layer with dimensions of 2x2 mm. This domain is discretized into a grid consisting of 100 x 100 LPs. Each LP represents a cell with an approximate diameter of 20 μm. Two scenarios are examined in this study, defined as *isotropic* and *gradient*. Both scenarios share the same set of parameters, except for the initial and boundary conditions in the glucose layer.

In the isotropic case, the initial glucose concentration was set to 5.5 mM over the entire domain (initial condition), mimicking a physiological level of glucose in the ECM [34], the concentration was fixed at the boundary of the domain (5.5 mM). As a result, the tumour spheroid placed in the centre of the domain is subjected to an isotropic chemical stimulus, analogous to what is qualitatively reported in **Figure 1a**. In the gradient case, we defined different boundary conditions at the left and right edges of the domain, imposing a fixed glucose concentrations of 8 [34,35] and 3 mM [36,37], respectively . As a consequence, the initial condition is a linear concentration gradient 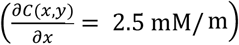, with an average concentration (and at the centre of the domain) still equal to 5.5 mM, as in the isotropic case. The initial concentration is constant along the y-axis.

Tumour spheroid is initially represented as an aggregate of few cells, occupying a sub-domain of radius 50 *μ*m (21 cells) in the centre of the lattice representing the ECM, which, for the sake of simplicity, is assumed to be composed of collagen. Spheroid growth was simulated for 48 h (simulated time). During the experiment spheroid evolves while individual cells proliferate and migrate invading ECM under the chemotactic stimuli of glucose concentration.

#### 2.1.2. Glucose layer

In the model, glucose is defined as a function of the spatial variable ***<u>x</u>*** = (x,y) and time *t*. Dynamics of glucose concentration field *C(****<u>x</u>***,*t)* in time and space is determined by solving the classical Fickian reaction-diffusion equation:

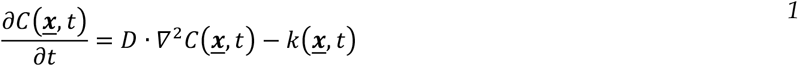

where, *D=*7 · 10^−10^ m^2^/s [38] is the diffusion coefficient of glucose in the collagen-based ECM and *k(****<u>x</u>***,*t)* is the consumption rate of glucose by the cells. Computational cost to obtain the numerical solution of **Equation 1** can be high, hence, given the large disparity between the time-scales of cell division (hours to days) and glucose diffusion (seconds), cellular proliferation has been treated as an adiabatic perturbation on the chemical field [29]. Thus, using the adiabatic perturbation approximation, **Equation 1** is approximated as a pseudo-stationary problem [39] :

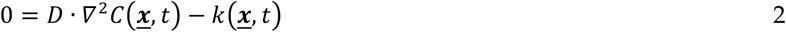

which is numerically solved by the method of simultaneous over-relaxation with Chebyshev acceleration [29]. This method requires to discretise the differential equation in terms of finite-differences, where *y* and *x* represent the row and column indices of the elements on the grid, and Δ is the size of a single LP (i.e., each of the two edges along *x* and *y*):

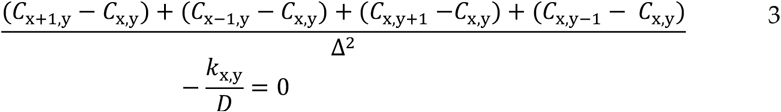

where Δ is the size of a single LP (in our case 20 *μ*m). **Equation 3** can be rearranged by defining its residual *ξ*_x,y_ as:

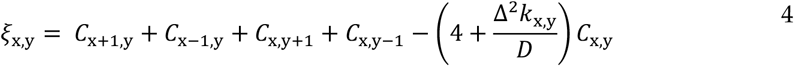

And calculating an approximate solution for the concentration field at the next simulation time step t + dt as:

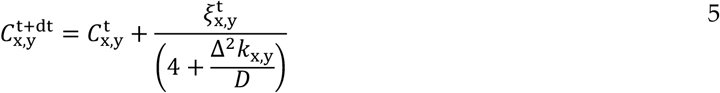

Glucose consumption by cells (in LPs where cells are present) is calculated according to a Michaelis-Menten kinetics:

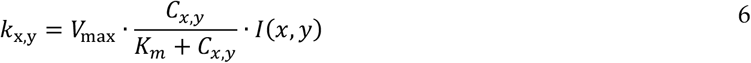

Where *V*_*max*_ = 0.05 mol/(m^3^ · s) [40] is the maximum consumption rate, *K*_*m*_ = 2 mol/m^3^ [40] is the Michaelis-Menten constant, and *I* is a Boolean indicator that takes on the value of 1 or 0 for lattice points that contain or not a cell. Because in our model two different cell phenotype (active and starved, see next section) able to consume glucose can occupy LPs, the indicator *I* in Equation 6 is calculated as the sum of two Dirac delta functions, one for each cell phenotype:

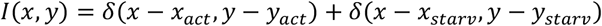

where (*x*_act_, *y*_act_) and (*x*_star_, *y*_*starv*_) are the coordinates of the LPs where active and starving cells are located. The use of the Dirac delta function allows for the representation of a point-like object, in this case the presence of a cell in a specific LP.

#### 2.1.3. Cellular layer and cell phenotypes

In the cellular layer of our model, the dynamics of individual cells are governed by specific rules that govern cellular dynamics and phenotyping, including migration, proliferation, starvation, and death.

Cell time evolution is governed by nutrient condition. In the cellular layer, three different cellular phenotypes are distinguished: active cells, starving (or quiescent) cells, and necrotic cells. The occurrence of various phenotypes depends on the cellular microenvironment and affect the metabolic activity of the cells. Two critical glucose concentration define the threshold for starving and necrotic cells (*C*_starv_ > *C*_nec_). A cell is active if the local glucose concentration *C*_x,y_ (i.e., concentration in the LP occupied) is above both these pre-defined thresholds, *C*_x,y_ > *C*_starv_ > *C*_nec_ . An active cell consumes glucose according to **Equation 6** and its metabolism include the possibility to migrate and eventually duplicates upon the completion of its cell cycle.

If *C*_nec_ < *C*_x,y_ < *C*_starv_, cell do not have enough nutrients to duplicate, and starves. It still consumes residual glucose and can migrate, looking for more nutrient-rich LPs. As time goes on, if *C*_x,y_ rises back above *C*_starv_, a starved cell can become active again, unless it remaines in the starved state for too long going in apoptosis (see next section).

If the nutrient concentration is further lower, *C*_x,y_ < *C*_nec_, cell undergoes necrosis. This process is irreversible, independently on any future change in glucose concentration and from that time on, necrotic cell cannot proliferate, migrate, or consume glucose.

#### 2.1.4. Rules governing cellular dynamics

The dynamic evolution of cells is highly dependent on nutrient availability, particularly glucose concentration. In response to nutrient availability, cells can undergo different fates, including necrosis and apoptosis (cell death).

Each non-necrotic cell in cellular layer can undergo migration or (if active) proliferation. The two mechanisms are regulated by two independent characteristic times, defined as migration time T_*m*_, and duplication time T_*d*_.

##### a) Cell migration

As simulation time goes on, every time interval T_*m*_, a cell can migrate. The probability of the cell to migrate is affected by cell-cell interactions if the cell under evaluation is attached to cell cluster and not isolated (detached). If a migration event occurs, cell will occupy one of its eight neighbouring LPs, provided the target location is empty (**Figure 1b**), alternatively, if migration does not occur, the cell remains in the same position until next T_*m*_.

When a cell tries to migrate, it must overcome cell adhesion defined by a probability *P*_adh_, defined as:

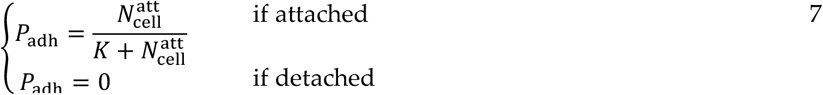

where 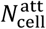 is the number of attached cells confining with the cell of interest 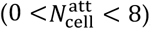, while *K* (K ∈ ℛ) is a parameter quantifying the role of cell-cell adhesion. For K=0 cell-cell adhesion is supposed to be so strong to inhibit the possibility to have any movement to the cells, while high values of K are related to a weak cell-cell interaction, and higher motility of the cells. When an attached cell attempts to migrate, a random number (0 < *σ* < 1) is generated and compared to *P*_adh_. If *σ* < *P*_adh_, the migration fails, until next T_*m*_. If *σ* > *P*_adh_ migration is allowed and a change in cell adhesion state can be induced; in our model, for simplicity, we assume that cell detachment is irreversible, i.e., once detached a cell will not be allowed to attach again. If the cell of interest is already detached (*P*_adh_ = 0), it shows typical behaviour of isolated cells and is expected to migrate every T_*m*_.

If migration is allowed, cell has to choose a motility direction, which is defined according to a biased random walk approach. Each of the nine possible positions *P*_*i*_ (*i* = 1, 2, …, 9) that a cell can take (**Figure 1b**), including the position already occupied (P_5_) is associated with an numeric interval *R*_*i*_ = [*a*_*i*−1_, *a*_*i*_ [. The 9 intervals are of different size (|*R*_*i*_ | = *a*_*i*_ − *a*_*i*−1_), are contiguous and non overlapping, and span the entire range 0-1 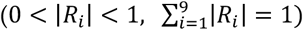 (**Figure 1c**). A further random number 0 ≤ *λ* ≤ 1 is generated and compared to the *R*_*i*_ intervals, if *λ* ∈ *R*_*i*_ the cell migrates towards the direction *P*_*i*_, if the corresponding LP is empty. In particular, if *λ* ∈ *R*_5_, i.e., the LP already occupied by the cell, cell does not migrate. If the cell, upon arrival at its new location, is not contiguous with any attached cells, it undergoes the transition to the detached state, provided it is not already in that state.

To evaluate the range of *R*_*i*_, and define probability of migration in each direction, a score *S*_*i*_ is evaluated for each of the 9 candidate positions as:

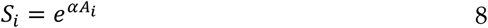

where *α* ∈ *ℛ* is a parameter describing glucose chemotactic sensitivity of the cell, and *A*_*i*_ is the local glucose specific gradient, defined as *A*_*i*_ = (*C*_*i*_ − *C*_5_)/*C*_5_. *S*_*i*_ values are further normalized 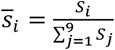. Finally, the ranges *R*_*i*_ = [*a*_*i*_, *a*_*i*+1_[are calculated as 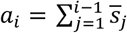 (**Figure 1c**).

##### b) Cell proliferation

As simulation time goes on, every time interval T_*d*_, active cells proliferate, generating a new daughter cell, identical to its progenitor. The new cell is located randomly in one of the free positions among the eight LPs surrounding the progenitor. If no empty spot is available, the new cell’s location is chosen identifying the direction where the minimum number of cells separates the progenitor from the edge of the cluster. All the cells along the selected direction shift one position away from the progenitor cell, and the new-born cell occupies the vacancy. It is worth mentioning our model takes memory of the history of each cell, to consider the possible cell dependent variables, such as random mutations of the cell parameters, even if in this work this feature is not used. In each time step all cells are considered the same, with the only difference being among the active, starved and necrotic phenotypes, while the cell dynamic evolution is dependent on the nutrient availability only. The code implemented in our model allows also for more complex interactions. Another simplification of our model is that only active cells attached to the spheroid can duplicate, while detached cells are expected to enter irreversibly in a migration state, unless nutrient availability induce their death.

##### c) Cell death

In our model, if a cell remains in starved state for a time longer than apoptosis time (T_apop_) it can spontaneously die. This biological event known as apoptosis [41] is a mechanism of defence of the cells to prevent propagation of lesions to the future generation. In our model apoptosis corresponds in degradation of the cell, which is dissolved in the ECM, and leaves the LP previously occupied as empty. It is worth saying that in case glucose concentration reduces further, down to values below *C*_nec_ while the cell is in already in starved state, cell does not go in apoptosis, but evolves in the necrotic state, where will remain indefinitely. Necrotic cells, according to our model, do not proliferate nor migrate, but continue indefinitely to occupy the same LP. The flow chart of the whole cell dynamic algorithm is reported in **Figure 2**. The model implementation of cell dynamic is also summarised as sequence of operation in figure caption.

**Figure 2.**
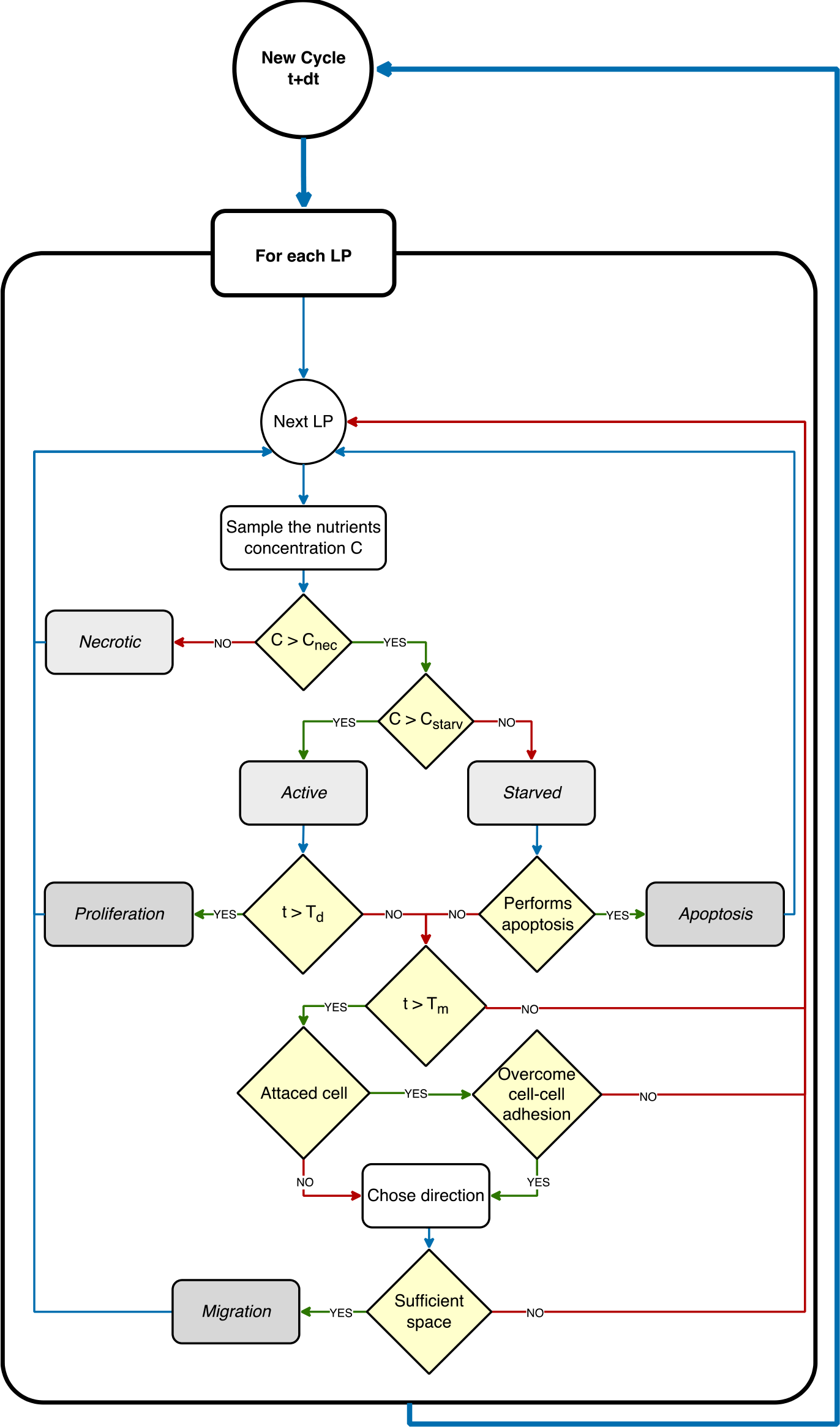
Cell dynamic simulation cycle. The CA model iteratively scans the Cellular Layer, where each LP represents a cell. If LP is occupied by a cell, the glucose concentration C_x,y_ is sampled from Glucose layer; if C_x,y_<C_nec_ the cell is classified as necrotic; if C_nec_ < C_x,y_ < C_starv_, the cell is classified as starved; if C_x,y_ > C_starv_, the cell is classified as active. Starved cells after T_apop_ undergo apoptosis and release the occupied LP. Active cells after T_d_ will proliferate. Active cells after T_m_ can migrate, according to migration rules.

#### 2.1.5. Analysis

In this study, we focused on analysing the effects of four key input parameters on the dynamic evolution of cell spheroids in our model: *T*_*d*_, *T*_*m*_, *K, α*. By investigating the influence of these parameters, we aimed to gain insights into the roles of cell adhesion, proliferation, and migration in tumour growth and invasion.

In **Table 1**, the output variables computed in our model are briefly summarised and they include the number of cells attached within the spheroid 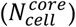, the number of cells that have migrated away from the spheroid 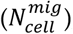, and the total number of cells 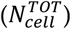. Additionally, the percentages of adhered and migrated cells relative to the total number were calculated as 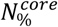 and 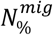, respectively. The ratio of adhered to migrated cells, *ϕ*, was also determined by calculating the ratio of 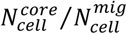, and considered as a measure of the invasiveness of the tumour. The lower is *ϕ*, the higher is the tendency of the cancer cells to invade and colonize the surrounding ECM.

**Table 1.**
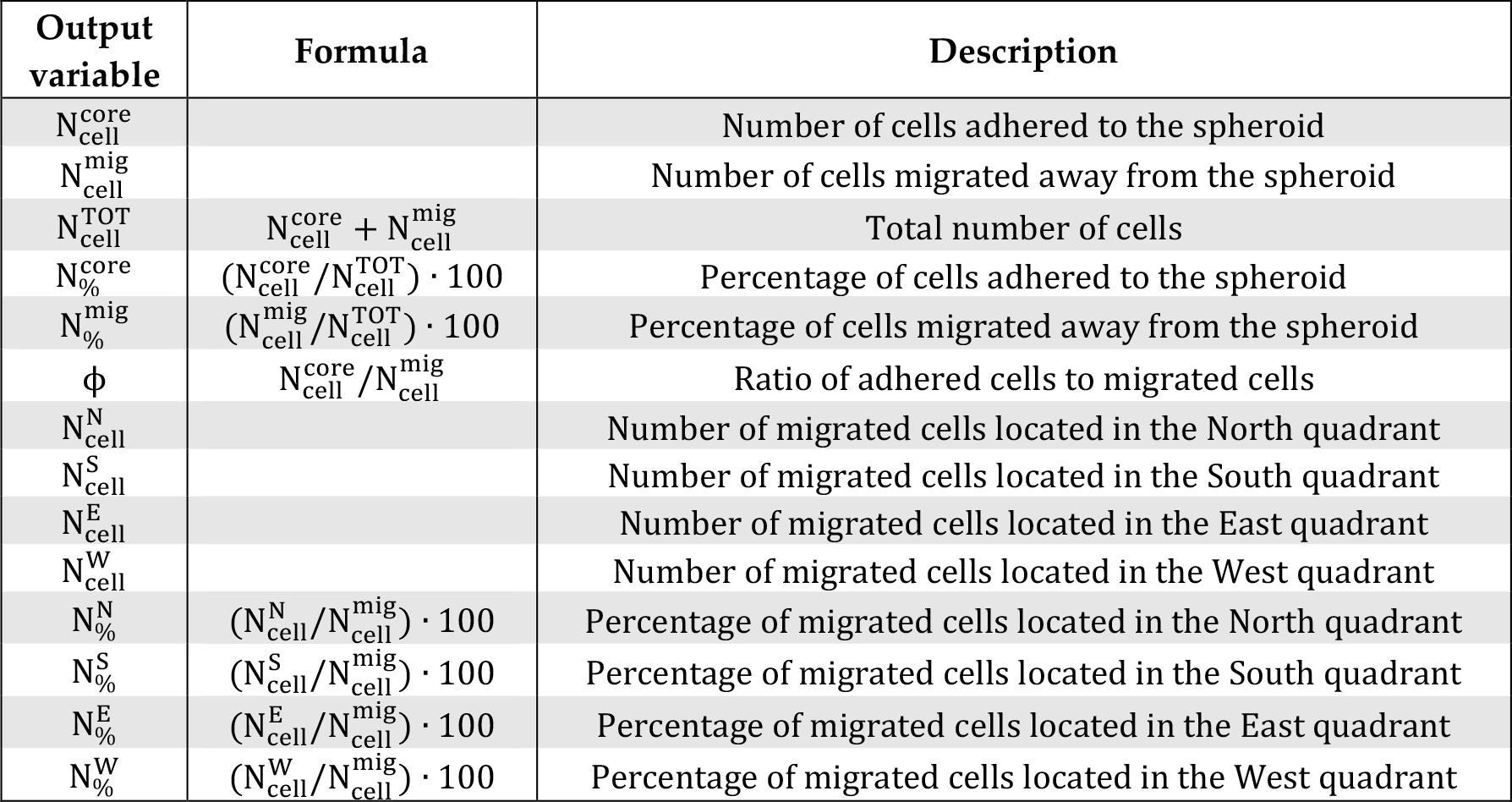
Model outputs.

Furthermore, the spatial domain in which the spheroid is located was divided into four sectors according to the diagonals of the square domain and named as North (N), South (S), East (E), and West (W) (**Figure S1**). The number of migrated cells located in each of these four quadrants were counted and represented as 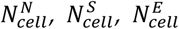, and 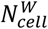. The corresponding percentages of migrated cells in each quadrant relative to the total number of migrated cells 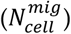 were also calculated.

### 2.2. Statistical Analysis

In this study, we employed statistical analysis techniques to gain a deeper understanding of our model by conducting both global and local sensitivity analyses. GSA enables us to assess the collective impact of perturbing all input parameters simultaneously, providing insights into how variations in multiple parameters influence the system as a whole, while LSA allows us to investigate the effects of perturbing individual parameters, providing detailed information on the sensitivity of the system to specific parameter changes. This approach allows us to identify the parameters that have a higher impact on specific aspect of the model, and to identify the controlling mechanism that drive the entire dynamic of the system. In our model, we investigated in particular the effect of varying the input parameters related to cell duplication time, cell migration, chemotactic motility and cell-cell adhesion: T_d_, T_m_, K, and *α*.

#### 2.2.1. Average and variance convergence

Since stochastic events and values of randomly generated numbers govern our HCA model, each simulation run is unique even if the values of parameters, initial conditions and boundary conditions are kept the same. For this reason, the outcomes of one single simulation for a given set of the four input parameters (T_d_, T_m_, K and *α*) is not enough to guarantee the statistical significance of the results. Therefore, for each set of input parameters, the simulation is replicated *n* times, defining the actual outcomes as the arithmetic average of the *n* iteration. To identify a value of *n* reliable from the statistical point of view, convergence analysis was performed to study the effect of number of simulation replicates on model outputs (see **Supplementary Materials, Figure S2**). Convergence was verified on 4 different set of model parameters, for the gradient experimental condition, running up to 100 simulations for each set. For *n* ≅ 10 all the model outcomes become constant and independent on *n* (data reported in **Figure S2**) for each of the 4 set of parameters investigated. Therefore, in the results presented below, each simulation was reiterated and mediated for *n* = 40.

#### 2.2.2. Global Sensitivity Analysis

To explore the impact of simultaneous parameter perturbations on key model outputs, a GSA was conducted, according to protocols reported in literature [42-44]. In brief, a Latin Hypercube Sampling (LHS) was used to sample the multidimensional parameter space generating a set of 500 combinations of the four key parameters of interest (α ∈ [0, 30], K ∈ [0, 4], T_d_∈ [10, 60 h] [45,46] and T_m_ ∈ [10, 120 min] [47]).

As there is no literature available to guide the choice of ranges for α and K, the selection was based on informed judgment and biological context. For the α parameter, which is an index of cell chemotaxis, the lower limit of 0 represents a situation where cells do not sense concentration gradients, while the positive range reflects a scenario where cells are attracted to increasing glucose concentrations. The upper limit of 30 was chosen to provide a degree of certainty that most cells would localize in regions that are rich in nutrients. For the K parameter, which is an index of cell-cell adhesion strength, the lower limit of 0 correspond to strongest cell adhesion which inhibit any cell movement, while the upper limit of 4 was chosen arbitrarily to represent an adhesion strength weak enough for all cells to detach from the spheroid in the temporal window of 48h.

For each of the 500 combinations of parameters a simulation of n = 10 iterations was run. A Multivariate Linear Regression Analysis (MLRA) was performed using the builtin MATLAB function *mvregress* on whole set of samples to obtain the linear regression coefficients for each parameter. The MLRA technique is based on the idea to express the output of interest *Y*_*i*_ as a linear function of the input parameters *X*_*j*_, as shown in **Equation 9**:

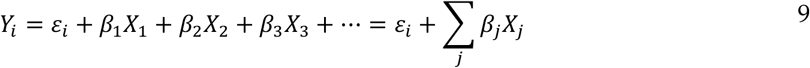

*ε*_*i*_ is an error term, and *β*_*j*_ are the regression coefficients of the input parameters. In our case, there are four input parameters *X*_*j*_ (T_d_, T_m_, K and *α*) and nine outputs *Y*_*i*_, which are 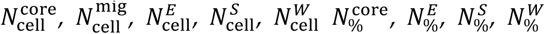. This subset of outputs *Y*_*i*_ was chosen to be linearly independent, as required by MLRA. The entire procedure (LHS and MLRA) was repeated 10 times, on ten different sets of 500 combinations of the input parameters *X*_*j*_ values combinations. As result four regression coefficient distributions were obtained. Each distribution (one for each input parameter *X*_*j*_) represent the measure of the sensitivity index (SI) of the respective parameter.

Finally, a one-way ANOVA was performed, followed by Tukey’s test, (test using the built-in MATLAB functions *anova1* and *multcompare*) on the SI distributions. This procedure allowed to rank the parameters according to their relative significance in affecting model outputs. The GSA was conducted for both the isotropic and gradient cases.

#### 2.2.3. Local Sensitivity Analysis

Each input parameter was perturbed independently while keeping constant the others at their respective baseline values. Parameters range chosen are the following: chemotactic index *α* [-30, 30] (basal value α = 6), cell-cell adhesion parameter K [0, 4] (basal value K = 0.2), doubling time T_d_ [10, 60h] (basal value T_d_ = 18h), migration time T_m_ [10, 120 min] (basal value T_m_ = 30min). For α, the range is set between -30 and 30 to account for a chemo-repellent effect. The baseline value of 6 was arbitrarily chosen based on trial and error in order to represent a moderate chemotactic force. For K, the range is still between 0 and 4, with a baseline value of 0.2. Again, the baseline value was selected through trial and error to ensure a balanced level of adhesion strength. For each parameter, 150 values were sampled uniformly and randomly within the specified range. Also in this case, the LSA was conducted for both the isotropic and gradient cases.

## 3. Results

In this section, we present the results of the numerical model simulations, which aimed to study the evolution of a tumour spheroid under two different glucose concentration profiles conditions, defined as isotopic and gradient. The results are organized into three main paragraphs. In the first paragraph, we present the results of the simulation of the tumour spheroid evolution assuming baseline values for the input parameters of the model. In the second paragraph, we report the result of a GSA used to identify the key parameters that have the most significant impact on the tumour evolution. Finally, in the third paragraph, we report the results of a LSA of the parameters to further understand the relationship between the parameters and the tumour evolution.

### 3.1. Baseline case

In this paragraph, we present the results of the simulation of the tumour spheroid evolution imposing baseline values to the four input parameters (i.e., α=6, K=0.2, T_d_=18 h and T_m_=30 min) both under isotropic and gradient conditions.

Figure 3. displays snapshots of the temporal evolution of the Cellular Layer and Glucose Layer at five time points (0h, 12h, 24h, 36h, 48h) for both isotropic (**Figure 3.a** and **b**) and gradient (**Figure 3.c** and **d**) conditions. The Cellular Layer (**Figure 3.a** and **c**) displays the extracellular matrix (ECM) in dark blue, with spheroid in the centre with attached and detached cells respectively reported in orange and light blue. The Glucose Layer (**Figure 3.b** and **d**) displays the concentration field of glucose in a colour scale, ranging from blue (low concentration) to red (high concentration).

The general behaviour observed is that the main tumour mass grows in times and consumes nutrients, inducing the formation of concentration gradients from the surrounding to the cell populated area. This chemical stress stimulates cell motion and leads cells detachment and migration toward regions less populated where higher concentration of nutrients is available in agreement with the predictions of diffusional instability theory [9], and its experimental verification [48]. The two conditions investigated present relevant differences. Under isotropic conditions, cells move radially away from the core, without any apparent preferential direction (**Figure 3.a**). Under gradient conditions, as consequence of the anisotropy in the stimulus, the detached cells tended to travel preferentially toward the source of nutrients at the left edge of the domain (**Figure3c**).

**Figure 3.**
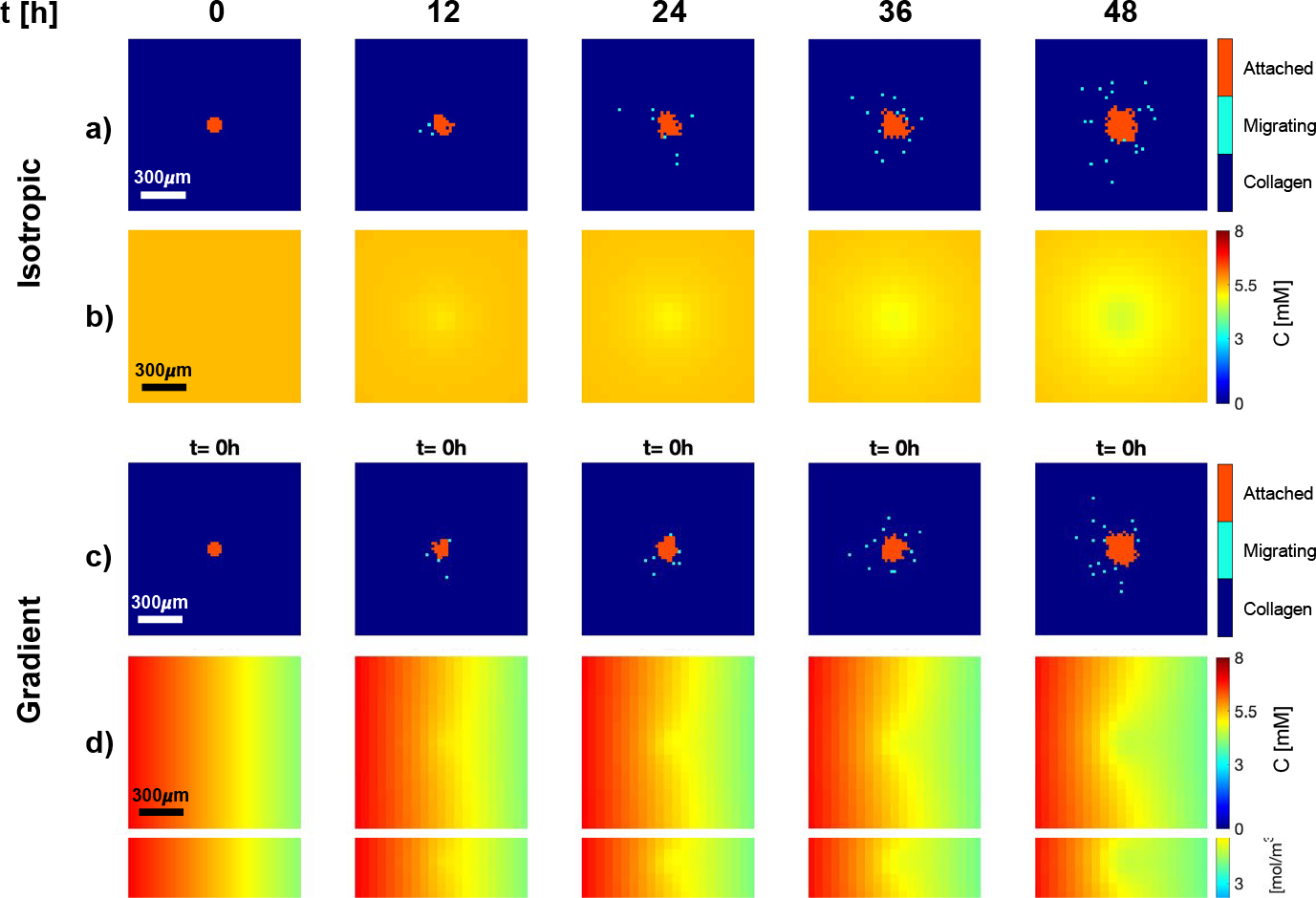
Baseline numerical solution. Snapshots of Cellular Layer (a,c) and Glucose Layer (b,d) at 0h, 12h, 24h, 36h, and 48h following seeding of cancerous cells in the centre of the simulation domain at 0h, under isotropic (a,b) and gradient (c,d) conditions. Cellular layers display cells attached to the core (orange), and cells detached from the core (light blue) which migrate in the collagen matrix (dark blue).

It is worth mentioning that, in the base values conditions here investigated, given the limited size of the spheroid and the short time frame simulated, the glucose concentration in the nutrient layer, determined by diffusion and consumption, remained always higher than *C*_starv_ = 0.1 mol/m^3^ . Therefore, all cells remained proliferative, and neither a necrotic core, nor a starvation rim were formed in the cellular layer. Running a simulation on a bigger domain (5 mm x 5 mm) and for a longer time (20 days) the appearance of necrotic cells was observed, as expected (results are reported in the supplementary, **Figure S3**).

To investigate the observed phenomenon quantitatively, and evaluate numerical measures of tumour evolution in time, we calculated the total number of cells 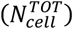 and the number of cells adhering to the spheroid 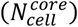 as a measure of tumor growth. The ratio of 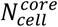 respect to cells migrated away from the spheroid 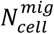, defined as *∅*, allows to compare proliferation and migration and it is a measure of the invasiveness of the tumour. We counted independently the number of cells migrated in the four sectors (**Figure S1**) 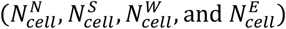, and the respective percentage 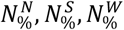, and 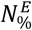). The comparison of these 4 values is a measurement of the anisotropy in cancer invasion, and directional response to chemotactic stimuli (diffusional instability).

Figure 4. presents the time-course plots of isotropic conditions and gradient conditions (**Figure 4.a-d** and **e-g**, in top and bottom part of the figure, respectively). The solid lines report the average output (*n* = 40) values, and the ribbons represent the associated standard error. The orange data reports the number (*n* = 40) of cells attached to the spheroid 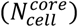, while the light blue data represent detached cells 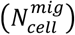. The **Figures 4.a** and **e** report the total cell count over time, while graphs in **Figures 4.b** and **f** report percentages of core and migrating cells. The **Figures 4.c** and **g** shows the time-course plot of the ratio *ϕ*.

The results show that the tumour core 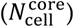 has an initial fast grow over time with an exponential trend for the first 24-36h, after this initial fast expansion, 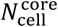 continues growing almost linearly (**Figure4a, e**). The number of cells detaching from the main body and migrating 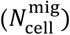 increases linearly over time, starting from the initial value of 0, for the entire temporal window investigated (**Figure4a, e**). Accordingly, the percentage of migrated cells 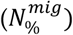 increases while the percentage of adhering cells 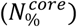 decreases as time goes on. After an initial period of instability, both the percentage of adherent and migrated cells reach a steady-state plateau, with the core cells 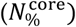 representing ∼80% of the total number of cells in the domain (**Figures 4b, f**). This steady state can be easily seen by the constant value of the ratio between the number of adherent cells and the migrated cells (*ϕ*=4), after 24h (**Figures 4c, g**). The result is due to the fact that both the size of the spheroid and the number of migrating cells increase linearly over time, with different rates, for long times. The system in other terms reaches a steady state where the generation of cells due to proliferation and the flow of cells leaving the spheroid are balanced. The above observed behaviour is comparable in terms of total cells in the core and migrating, among the two conditions here investigated, i.e., isotropic and gradient, while a relevant difference can be observed in the direction of migration of the invading cells.

**Figure 4.**
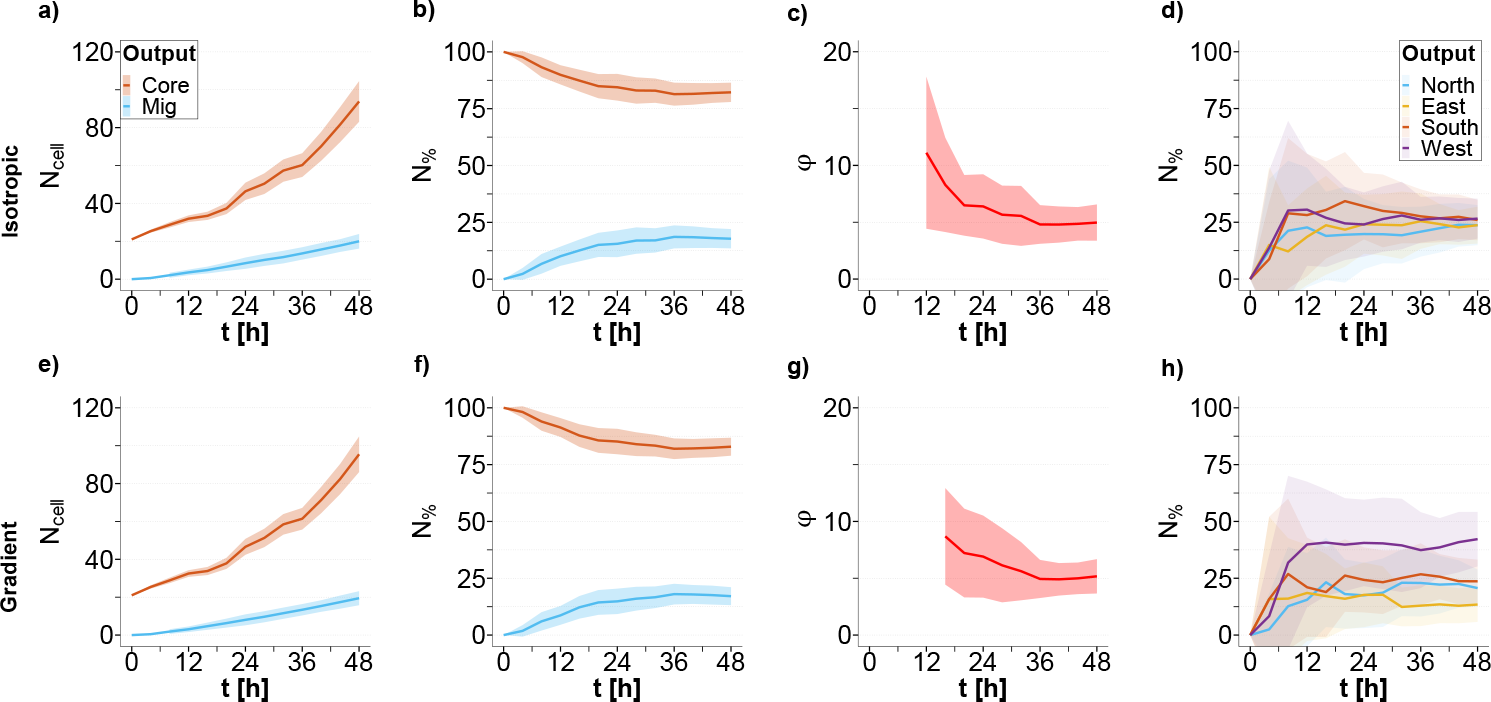
Quantification of baseline solution. Results of the simulation assuming basal conditions with input parameters *α*=6; *K*=0.2, *TT*=18*h, TT*=30*Tin* in isotropic (a-d) and gradient case (e-h); **a**,**e)** core cell number (orange) and migrated cell number in time (light blue); **b**,**f)** core cell percentage (orange), migrated cell percentage (light blue); **c**,**g)** migrated cells number ratio *ϕ*; **d**,**h)** percentage of cells migrated in the four directions North (light blue), East (yellow), South (orange) and West (purple) in time. The solid lines represent the means (n=40), and the ribbons represents the standard errors.

**Figures 4.d** and **h** reports the percentages of the migrated cells in the four different directions, defined according to the source of nutrients (see **Figure S1**). In isotropic conditions the detached cells do not show any preferential migratory direction and they uniformly distribute into the four sectors of the domain 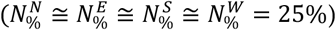 (**Figure4.d**). On the other hand, in the gradient case, a significant difference (P<0.005) in the percentage of detached cells that migrate in the four directions is observed. As shown in **Figure 4.h**, nutrient gradient induces cells to move in a biased random walkway, such that a greater percentage of the population migrated towards the west end (higher glucose concentration). At the end of the simulation (t=48h) under anisotropic conditions, 42±12% of cells migrate towards the direction of increasing concentration gradient, and only 13±8% of cells move against the gradient, heading towards the East. The remaining migrating cells distribute almost uniformly between the remaining North and South sectors, with 21±8% and 24±10% of cells present in each sector respectively.

### 3.2. Global and sensitivity analysis (GSA)

To perform the analysis, we estimate the SI of each input parameter (*X*_*j*_) and rank them according to their importance.

In **Figure 5**, the regression coefficients obtained from MLRA for the isotropic and gradient cases are plotted. Each bar represents the regression coefficient mean, which is a measure of parameter SI. The higher SI, the greater is the sensitivity of the parameter in affecting a given model output. To determine the ranking of the relative sensitivity of model parameters, we conducted one-way ANOVA and post-hoc analysis (Tukey’s test) after checking that the regression coefficients follow a normal distribution, a crucial requirement for the two tests. The bars marked with an asterisk indicate the statistically significant parameters for each model output, obtained from MLRA.

**Figure 5.**
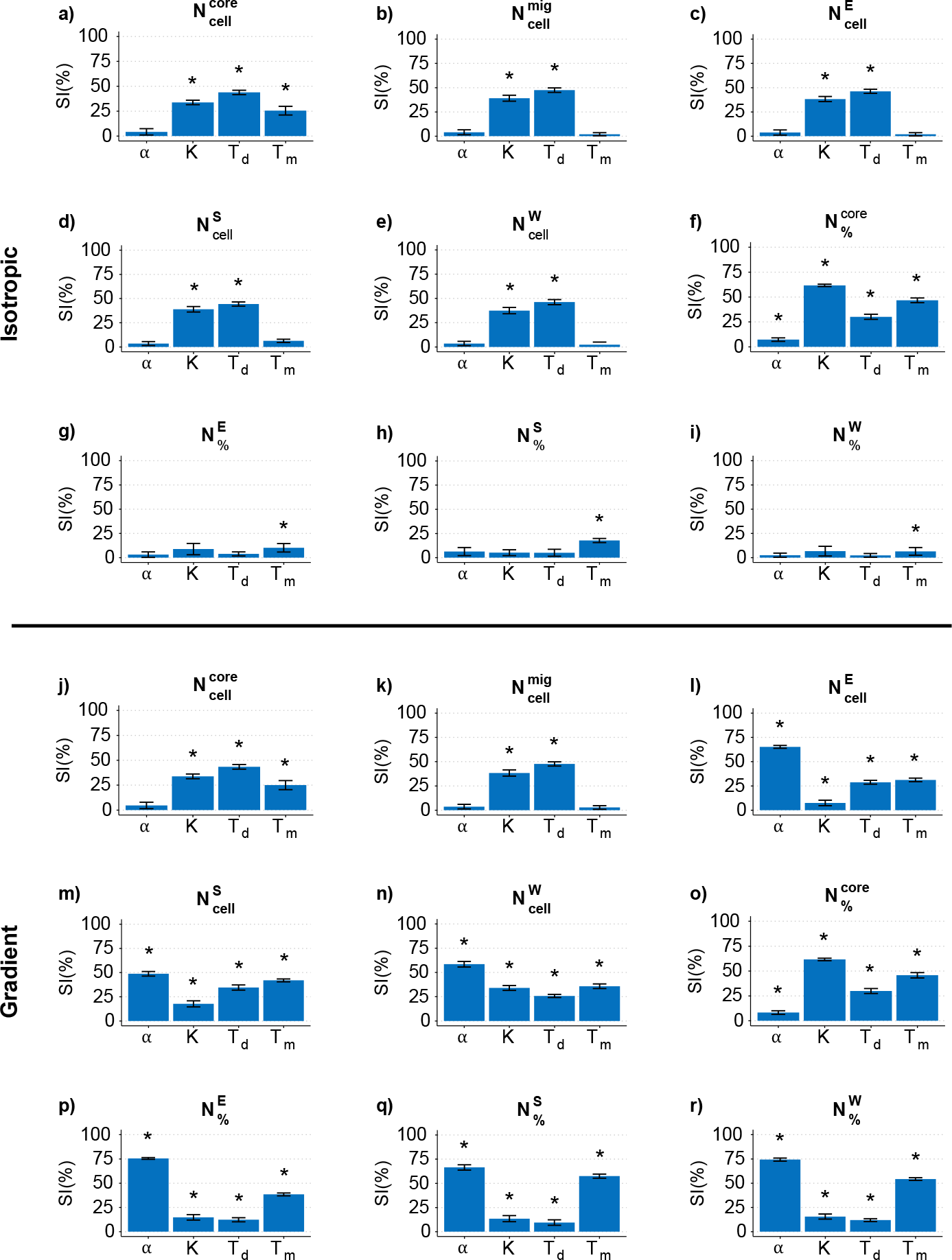
MLRA regression coefficients (SI) in isotropic case (a-i) and gradient case (j-r). The columns refer to the key parameter *α, K, T*_*d*_, *T*_*m*_ . The rows refer to the output of interest: **a**,**j)** core cell number; **b**,**k)** migrated cell number; **c**,**l)** number of cells migrated toward East; **d**,**m)** number of cells migrated toward South; **e**,**n)** number of cells migrated toward West; **f**,**o)** core cell percentage; **g**,**p)** percentage of cells migrated toward East; **h**,**q)** percentage of cells migrated toward South; **i**,**r)** percentage of cells migrated toward West. The asterisk indicates which parameter is significance (p<0.05) for the related output.

From the GSA, we observed that the 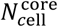 is independent on *α* (analogous result would be obtained measuring core area), but it is significantly affected by *T*_*d*_ (**Figure 5.a and j**). Number of migrated cells 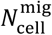 is independent on *α* and *T* under both isotropic and gradient conditions (see **Figure 5.b** and **k**). Looking at the migration direction-related outputs 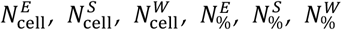, relevant differences are visible comparing the two conditions investigated. In the gradient case, all the migration direction-related outputs depend on all the four parameters, and in particular on *α* (**Figure 5.l-r**). This is expected since *α* is related to the chemotaxis. On the other hand, in the isotropic case, 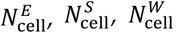, are dependent only on T_d_ and K (**Figure 5.c-e**), with the former being the most relevant, while 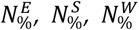 are affected only by *T*_*m*_ (**Figure 5.g-i**). From these observations, we see that: *α* influences only outputs related to the directionality of cell migration solely in the presence of a gradient, while it has no effect in isotropic conditions; the parameter K and T_d_ affect both the size of the spheroid and the number of cells detaching from it, in both isotropic and gradient conditions; Tm has a smaller effect on spheroid growth compared to K and Td but it significantly influences the directionality of cell migration only in the presence of a gradient.

### 3.3. Local sensitivity analysis (LSA)

In order to define the empirical functional relationship between parameters and model outputs, we conducted an LSA. The results of the LSA provide a deeper insight into the non-linear relationships between the parameters and the tumour evolution and highlight the regions of the parameter space where the response variables are most sensitive to changes.

We analysed the effect of the key input parameters *α, K, T*_*d*_, and *T*_*m*_ on all model outputs (see **Table 1**). However, to the sake of brevity, we are limiting our discussion here only to the most significant input-output relationship suggested by to the results we obtained from GSA.

#### 3.3.1. Chemotaxis sensitivity index *α*

As described in the materials and method section, the parameter *α* is an indicator of cell sensitivity to chemical stimuli, represented by glucose in this work. The higher the absolute value of *α*, the greater is the tendency of the cells to migrate following a given concentration gradient.

**Figure 6a** and **b** depict the fraction of migrating cells invading from the spheroid along the four sectors (north, south, west, and east, see **Figure S1**) of the domain, that are subject to different chemical stimuli, due to the geometry of our domain. Cells are counted after 48h of real time simulation and reported on the y-axis as the fraction of cells in each direction, the x-axis shows the *α* values. The solid lines represent the mean fraction of cells, while the ribbons represent the associated standard deviation calculated on the n = 40 iterations (see 2.2.1).

**Figure 6.**
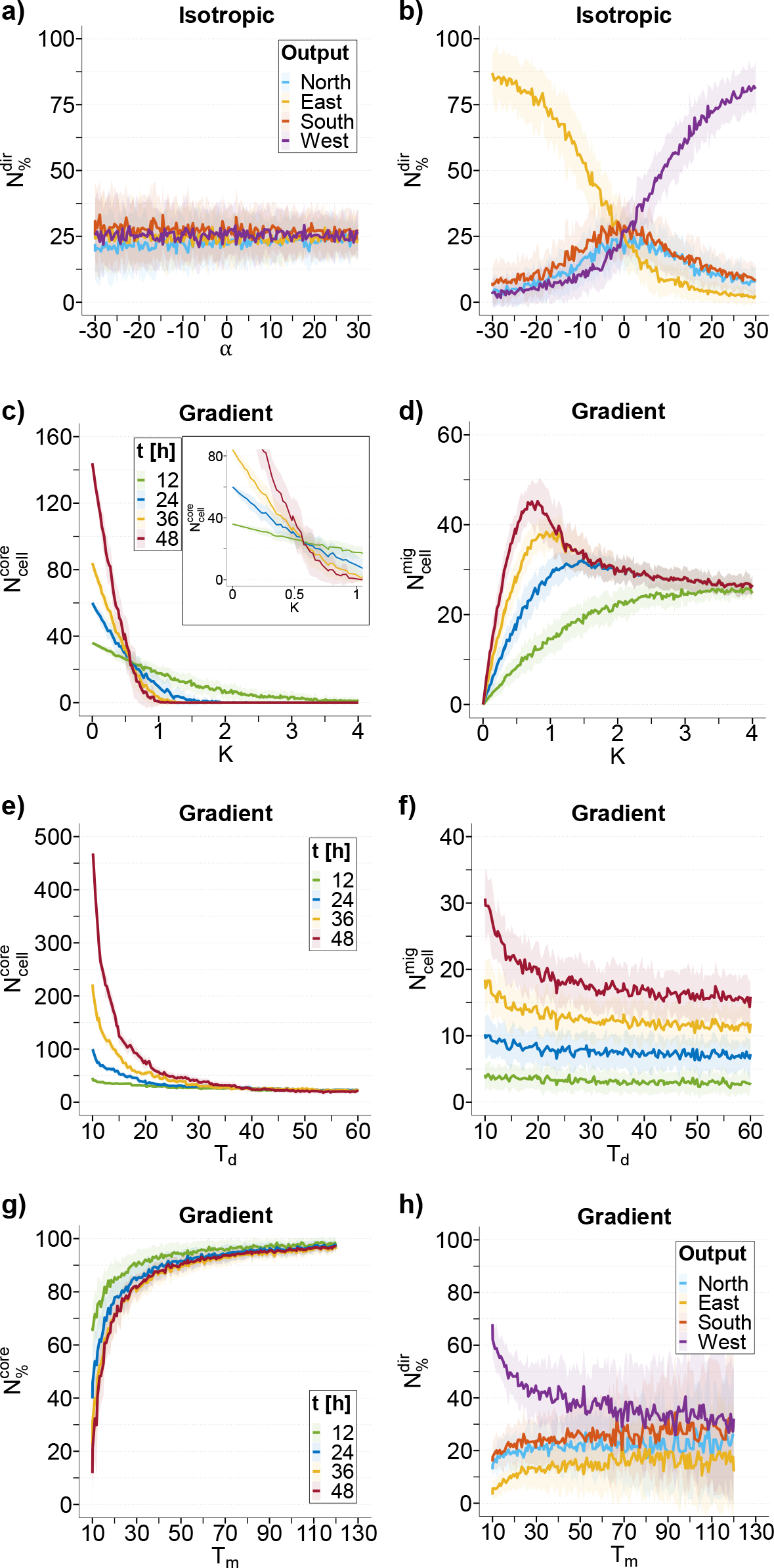
LSA most relevant results in according to the GSA. Results of LSA perturbing the input parameter *α*: **a)** percentage of cells migrated in the four directions North (light blue), East (yellow), South (orange) and West (purple) at 48h in isotropic case; **b)** percentage of cells migrated in the four directions North (light blue), East (yellow), South (orange) and West (purple) at 48h in gradient case. Results of LSA perturbing the input parameter *K*: **c)** core cell number at 12h (green), 24h (blue), 36h (yellow), 48h (red) in gradient case; **d)** migrated cell number at 12h (green), 24h (blue), 36h (yellow), 48h (red) in gradient case. Results of LSA perturbing the input parameter *T*d: **e)** core cell number at 12h (green), 24h (blue), 36h (yellow), 48h (red); **f)** migrated cell number at 12h (green), 24h (blue), 36h (yellow), 48h (red). Results of LSA perturbing the input parameter *T*m: **g)** core percentage at 12h (green), 24h (blue), 36h (yellow), 48h (red); **h)** percentage of cells migrated in the four directions North (light blue), East (yellow), South (orange) and West (purple) at 48h. The solid lines represent the means (n=40), and the ribbons represents the standard deviation.

Under isotropic conditions (**Figure 6.a**), the fraction of cells migrating in the four sectors is independent on the *α* parameter, with cells uniformly distributed throughout the entire domain 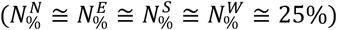 in agreement with the isotropic stimulus imposed.

In contrast, under anisotropic conditions (**Figure 6.b**), the distribution of migrating cells in the four sectors strongly depends on *α*. We investigated also negative values of *α* to study the potential impact of a chemorepellent (such as a catabolite, or a toxic drug) on the migration of cancerous cells. Three cases are distinguishable in the graph: *α* = 0, *α* > 0, and *α* < 0.

For *α* = 0, cells are insensitive to the concentration gradient, and the system restores a pseudo-isotropic condition, with migrating cells uniformly distributed 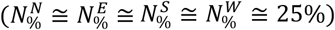, as in the case of **Figure 6.a**.

For *α* > 0, cells show a preferential migration towards the source of chemoattractant, that is West direction in our domain. As *α* increases, 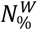 (purple) increases linearly up to reach a saturation for high values of *α* (*α* > 20), with the fraction of cells plateauing at around 85%. The fraction of cells in the remaining sectors decreases accordingly. It can be seen that 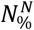 (light blue) and 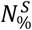 (orange) curves almost overlap, due to the North/South symmetry of our setup, while Est direction (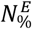, yellow) decreases more rapidly being opposite to the source of chemoattractant.

A completely inverted situation is found for *α* < 0, where cells move away from the source of chemical and show a preferential migration towards decreasing gradients, with directionalities of cells inverted. The trends of the curves associated with cells migrating towards the west and east are inverted but the overall trend of the chart is symmetric.

#### 3.3.2. Cell-cell adhesion parameter K

As described in the materials and method section, parameter *K* models cell-cell adhesion, the higher the value of *K*, the greater is the tendency of an attached cell to move to one of the adjacent positions. This implies a greater chance that a cell could detach from the spheroid and invade the surrounding. Low values of K are related to a limited tendency to invade.

As described in the GSA, parameter *K* mostly affects the number of cells connected together within the spheroid 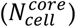, the number of detached cells that migrated away 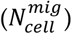, and their percentages (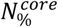 and 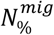). The general result being, as expected, that 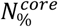 monotonically decreases as time goes on and K increases (data not showed).

The trends of 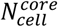 and 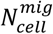 were evaluated at four different time points (12h, 24h, 36h, and 48h) as a function of the parameter *K* and are shown in **Figure 6.c** and **d** respectively. We report here data only for the gradient case, the results obtained were independent of the concentration field, as confirmed by the GSA. As in the previous paragraph, solid lines represent the mean value of cell number, while ribbons represent the associated standard deviation.

For simplicity, we can distinguish four scenarios: *K* = 0, 0 < *K* < 0.5, *K* = 0.5, and *K* > 0.5. When *K* = 0, cell adhesion is indissoluble, and so, 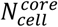 monotonically grows over time, while 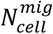 remains at 0. For values of 0 < *K* < 0.5, cell adhesion is still quite strong, and 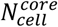 can still grow over time, although at a decreasing rate as K increases. This growth is slowed by the progressive detachment of migratory cells, which increases over time and as K increases.

For the critical value of *K* = 0.5, a steady-state equilibrium is established (**Figure 6.c, inset**). The number of cells adhering to the spheroid remains constant over time and is approximately equal to the initial number of cells (21 cells, t=0). On the other hand, the number of migratory cells evaluated at 48h (red) reaches its absolute maximum for this value of K. This indicates that there is a balance between the “flow” of cells leaving the tumour and the generation of new cells within the tumour.

For values of *K* > 0.5, cell adhesion weakens further, and the generation of new cells cannot compensate the outflow of migratory cells. The number of adhering cells monotonically decreases over time until the spheroid completely disintegrates at 48h. Similarly, 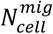 decreases from the peak reached for *K* = 0.5 until it reaches a plateau, as there is no longer a tumour core able to generate new cells (**Figure 6.d**). The higher the value of K (above 1), the earlier the time the tumour core disappears.

#### 3.3.3. Doubling time T_d_

Based on the GSA results, T_d_ mostly impacts the number of adherent 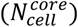 and migrating cells 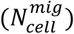. **Figure 6e** and **f** display the trends of 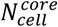 and 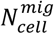, evaluated at four different time points (12h, 24h, 36h, 48h) as T_d_ variates. Again, we present the results only for the gradient case. Upon increasing *T*_*d*_, 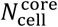 count (**Figure 6.e**) rapidly decreases, and a similar but less marked trend is followed by 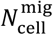 (**Figure 6.f**), confirming the results of the GSA again.

#### 3.3.4. Migration time T_m_

Once a cell is detached, *T*_*m*_ represents the time required by the cell to travel a distance equal to its diameter, thereby indicates motility of cells. Alternatively, when a cell is attached, its capacity to move is limited, and *T*_*m*_ represents the inverse of the frequency at which the attached cell attempts to move.

As described in the GSA, the parameter *T*_*m*_ mostly affects the fraction of cells comprising the spheroid 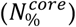 both in isotropic and anisotropic case, and only in the gradient case the fraction of migrating cells located in the four sectors 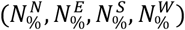.

**Figure 6.g** depict 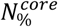 evaluated at four time points (12h, 24h, 36h, and 48h) in gradient conditions. Decreasing *T*_*m*_, the core reduces in size because a larger number of cells migrate toward the surroundings (**Figure 6.g**) due to an increased tendency of migration.

**Figure 6.h** shows 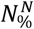 (light blue), 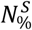 (orange), 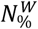 (purple), and 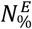 (yellow) evaluated at 48h as a function of T_m_. In the gradient case, the effect of *T*_*m*_ is observed by an increase in the cell population percentage migrating towards West (higher glucose concentration) when *T*_*m*_ decreases (**Figure 6h**). However, when *T*_*m*_ becomes sufficiently large, the four percentages 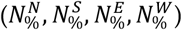 seem to converge to ∼25% value, which can be attributed to the reduced frequency of all cells to migrate.

## 4. Discussion

The HCA model developed in this study successfully simulated the temporal and spatial evolution of a tumour spheroid under different chemical stimuli. The model incorporates the key biological mechanisms driving tumour progression, such as cell proliferation, migration, apoptosis, and interactions with the extracellular matrix and neighbouring cells. In this study, the growth and invasiveness of an avascular tumour spheroid were simulated in two different chemical fields, isotropic and anisotropic, for 48 hours to study its evolution over time. For a fixed set of input parameters, and independently by the type of the chemical field, the spheroid core area increased over time, and a fraction of cells detached and invaded the tumour’s surroundings. The effect of an external glucose gradient was to convert cell migration from a random walk to a chemotaxis-driven biased random walk.

Through GSA and LSA, we were able to understand how the selected input parameters affect the system, including how the parameters interact with each other and how they act individually. The GSA revealed that impact of *α* was negligible in isotropic conditions but was a fundamental factor in governing tumour invasion under the gradient case; the parameters Td and K were critical for the evolution of the core area and the number of cells detaching from the spheroid core, in both chemical fields, while Tm has a similar but less effective influence on tumour growth and invasiveness.

The LSA quantitatively assessed the effect of parameter perturbation, showing that the value of *α* influenced the migration direction of cells towards or away from a positive gradient, while *α* equal to zero resulted in cells becoming insensitive to the gradient. The parameter K, representing cell-cell adhesion, played a crucial role in tumour evolution, with K=0.5 marking the border between scenarios where adhesion promoted proliferation, or the flux of migrating cells dissolved the tumour over time. Perturbing T_d_ led to a drastic decrease in the number of core cells and invading cells, while perturbing T_m_ had the opposite effect, accelerating tumour growth.

Despite its simplicity, the HCA model proved to be highly effective in simulating tumour growth and invasion over time, capturing essential features of tumour biology and reproducing realistic spatiotemporal patterns under different experimental conditions. The model’s versatility allows for easy modification and adaptation to include additional biological, chemical, and physical processes relevant to tumour growth and invasion. The model can be calibrated and validated using experimental data obtained from *in vitro* or in *vivo assays*, thus providing a more accurate representation of the biological system under investigation.

Moreover, the model can be used to design and predict the outcomes of new experiments setups, thereby reducing the need for extensive and costly experiments. By incorporating new biological parameters extracted from simple and rapid experiments, the model can simulate more complex scenarios beyond the limitations of laboratory-based assays. Future improvements can involve the integration of complex signalling pathways, immune responses, hypoxia, nutrient transport, and mechanical stresses, such as oncotic pressure or laminar flows affecting the tumour surface.

In conclusion, the HCA model represents a promising tool for investigating the mechanisms of tumour growth and invasion, as well as for guiding the design and assessment of novel therapeutic strategies. Its ability to capture the complexity of tumour biology and its adaptability to various experimental settings make it a valuable asset in cancer research.

## 5. Conclusions

This work introduces model based on the HCA (Hybrid Cellular Automaton) approach, capable of simulating tumour growth and cancer cell invasiveness. Model’s performance was tested, simulating the evolution of a 3D tumoral model, spheroid, at the early stages of its development, and under isotropic and anisotropic glucose concentration fields. Furthermore, this study investigated how perturbations in selected cellular parameters, including chemotactic sensitivity, cellular adhesion, doubling time, and cell motility, affected tumour dynamics.

These parameters have been ranked according to their influence on model outcomes by performing a GSA. The results demonstrated that the automata-based model accurately described the initial exponential-like growth of tumours and depicted cells adapting their migration mechanism in response to external chemical stimuli. The chemotactic index was identified as crucial for cellular migration in the presence of a chemoattractant gradient, but it was irrelevant for the tumour development in isotropic chemical fields. Cellular adhesion plays an essential role in tumour growth and metastasis. The study revealed an optimal value of parameter K, representing cellular adhesion, where the tumour produced the highest quantity of migrating cells while maintaining a constant size over time. Additionally, the investigation highlighted that a hypothetical cancer cell line characterized by high motility and a short doubling time generates cancers with fast growth and high invasiveness.

Future developments of this work involve the calibration and validation of the presented model using experimental data obtained from spheroid growth *in vitro*, enhancing its applicability and contribute to a better understanding of tumour behaviour.

## Supplementary Materials

The following supporting information can be downloaded at: www.mdpi.com/xxx/s1, Figure S1: Schematic illustration of the division into four sectors of the cellular layer; Figure S2: Convergence Analysis; Figure S3: Simulating Advanced Tumour Morphology and Temporal Changes under Isotropic Conditions.

## Author Contributions

Conceptualization, S.C. and P.D.; methodology, L.M., M.J.P. and P.D.; formal analysis, P.D.; investigation, L.M.; data curation, R.F. and L.M.; writing—original draft preparation, L.M. and R.F.; writing—review and editing, P.D., Z.W., V.C. and S.C.; supervision, S.C. and P.D.. All authors have read and agreed to the published version of the manuscript.

## Informed Consent Statement

Not applicable

## Data Availability Statement

The data presented in this study are available on request from the corresponding author.

### Acknowledgments

This work was partially supported by funding from the Cockrell Foundation (PD). This study was conducted under the umbrella of the International Academic Affiliation Agreement between Houston Methodist Academic Institute (Houston, TX, USA) and University of Naples Federico II (Napoli, Italy).

## Disclaimer/Publisher’s Note

The statements, opinions and data contained in all publications are solely those of the individual author(s) and contributor(s) and not of MDPI and/or the editor(s). MDPI and/or the editor(s) disclaim responsibility for any injury to people or property resulting from any ideas, methods, instructions or products referred to in the content.

