## Supplementary text for "Hybrid Cellular Automata Modelling Reveals the Effects of Glucose Gradients on Tumour Spheroid Growth"

*Correspondence should be addressed to:

#### *S1: Schematic illustration of the division into four sectors of the cellular layer*

The image depicts a representation of the cellular layer domain under the two analyzed conditions: isotropic and gradient. In both conditions, the tumor spheroid (depicted as an orange circle) is situated in the center of the domains which are divided into four sectors (North, South, West, East). The key distinction between the domains under the two conditions is the glucose concentration profile, depicted using a color gradient ranging from low (blue) to high (red) concentration. In the isotropic case, glucose concentration is radial, with maximum values along the domain edges, while the central region is nutrient-deficient due to assimilation by the central tumor. Consequently, all four sectors exhibit identical concentration profiles. In the gradient case, glucose concentration follows a linear profile, with the highest concentration at the left edge (West sector) and the lowest concentration at the right edge (East sector). The South and North sectors display symmetric nutrient concentrations with an intermediate concentration level (green).

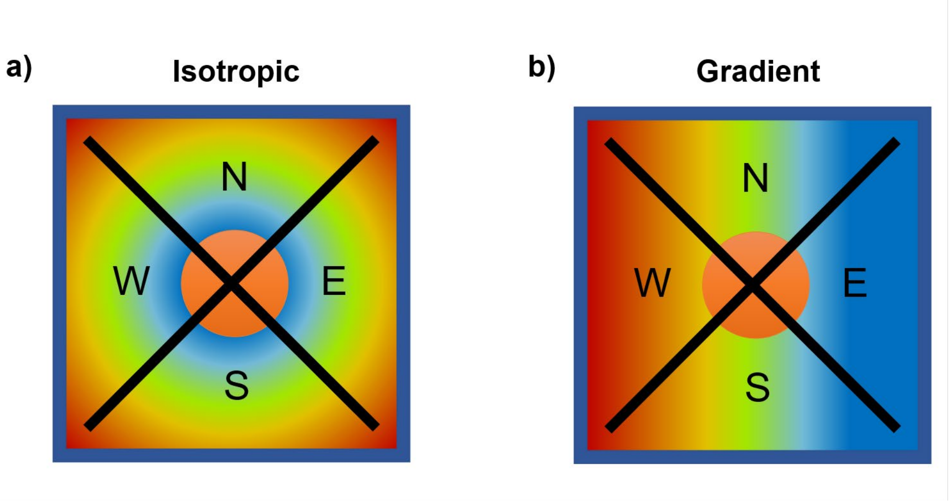


***Figure S1.*** *Qualitative representation of the domain in which is located the spheroid (orange circle) divided in four sectors (North, East, South, West) in isotropic (a) and gradient (b) conditions. The cellular layer and glucose layer has been superimposed.*

### *S2. Convergence Analysis*

Convergence analysis was performed to assess the stability and reliability of the simulations. Each simulation yielded slightly different results despite having the same parameters, initial conditions, and boundary conditions. To obtain statistically valid measurements, four sets of parameters were selected. For each set, 100 simulation iterations were performed. The trend of the mean and standard error of the simulations was examined as the number of iterations increased. The minimum number of iterations, n=10, was determined to ensure stability in terms of mean values and low standard error. To ensure robustness and account for variability, all subsequent simulations were iterated 40 times, unless explicitly stated otherwise.

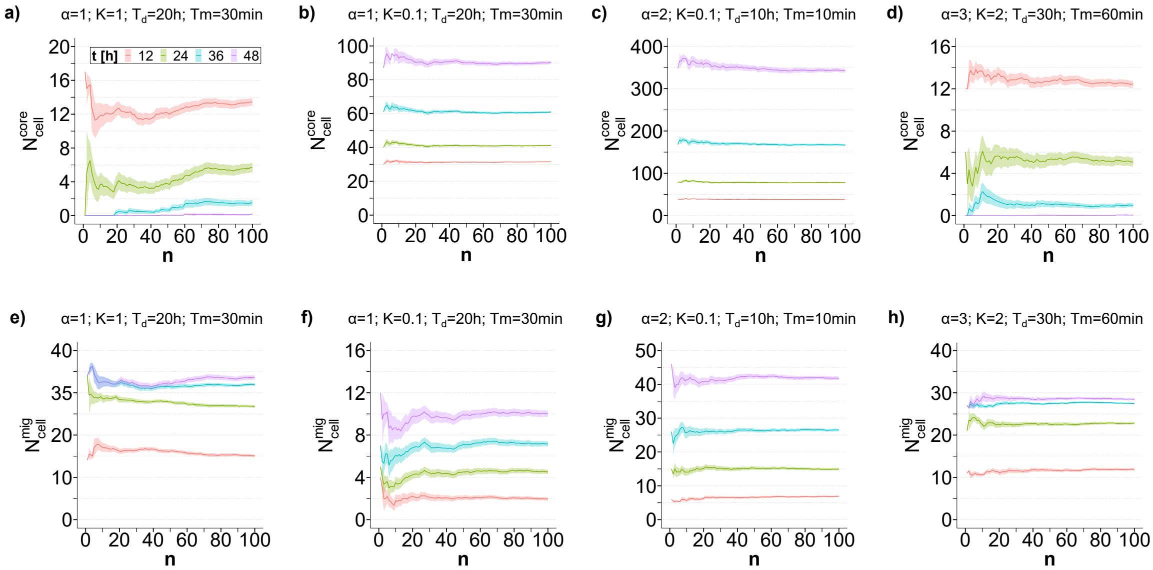


***Figure S2.*** *Convergence Analysis results computed assuming 4 randomic combinations of input parameters and evaluated at 4 time points (12h, 24h, 36h, and 48h).* ***a-d)*** *average number of cells attached to the spheroid core as function of the number of the simulation iterations n;* ***e-h)*** *average number of migrated cells as function of the number of the simulation iterations n. The solid lines represent the means (over the n iterations), and the ribbons represents the standard error.*

#### *S3. Simulating Advanced Tumour Morphology and Temporal Changes under Isotropic Conditions*

The scenario studied in the main article does not exhibit the formation of necrotic cores due to the limited size of the spheroid and the simulated time window, which hindered substantial tumor growth. However, this specific simulation was intentionally designed to highlight and demonstrate the accurate reproduction of advanced tumour morphology by the model. As expected, the simulation captures the presence of an outer rim comprising active and proliferative cells, an intermediate rim consisting of quiescent cells, and a central core of necrotic cells. Furthermore, the figure showcases the temporal changes in nuclear morphology, revealing its transformation from a circular conformation to the formation of lobes and protrusions towards nutrient-rich regions. These observed morphological alterations correspond to the predictions of the diffusional instability model.

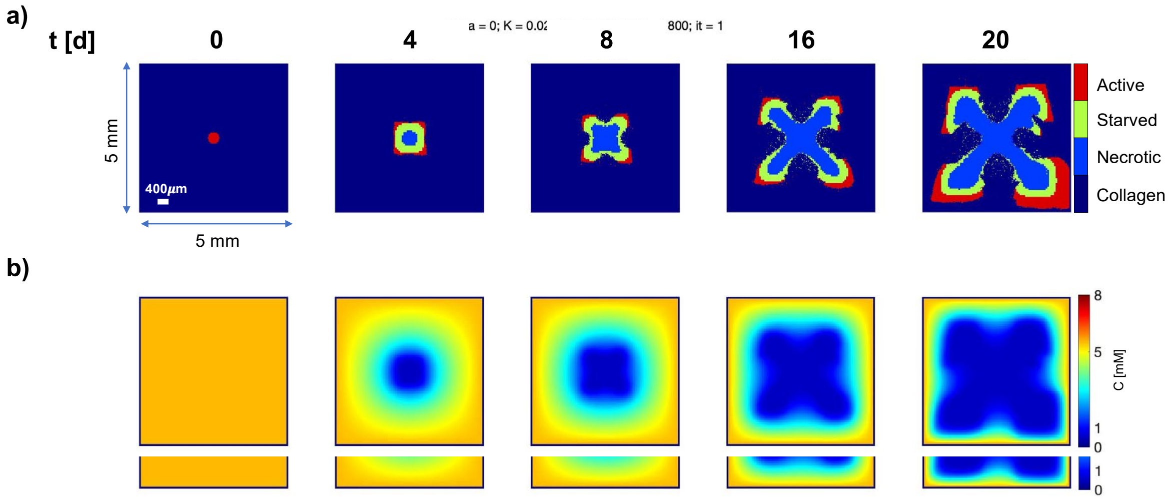


*Figure S3. Baseline numerical solution. Snapshots of Cellular Layer (a) and Glucose Layer (b) at 0, 4, 8, 16, and 20 days, following seeding of a tumour spheroid (diameter = 200μm) in the centre of the simulation domain (size 5mm x 5mm) at 0 day, under isotropic condition. Cellular layers display active (red), starved (green) and necrotic (blue) cells growing and evolving in a collagen matrix (dark blue).*
